# A common framework for identifying linkage rules across different types of interactions

**DOI:** 10.1101/024315

**Authors:** I. Bartomeus, D. Gravel, J.M. Tylianakis, M.A. Aizen, I. A. Dickie, M. Bernard-Verdier

## Abstract

Species interactions, ranging from antagonisms to mutualisms, form the architecture of biodiversity and determine ecosystem functioning. Understanding the rules responsible for who interacts with whom, as well as the functional consequences of these interspecific interactions, is central to predicting community dynamics and stability. Species traits *sensu lato* may affect different ecological processes determining species interactions through a two-step process. First, ecological and life-history traits govern species distributions and abundance, and hence determine species co-occurrence, which is a prerequisite for them to interact. Second, morphological traits between co-occurring potential interaction partners should match for the realization of an interaction. Moreover, inferring functioning from a network of interactions may require the incorporation of interaction efficiency. This efficiency may be also trait-mediated, and can depend on the extent of matching, or on morphological, physiological or behavioural traits. It has been shown that both neutral and trait-based models can predict the general structure of networks, but they rarely accurately predict individual interactions, suggesting that these models may be predicting the right structure for the wrong reason. We propose to move away from testing null models with a framework that explicitly models the probability of interaction among individuals given their traits. The proposed models integrate both neutral and trait-matching constraints while using only information about known interactions, thereby overcoming problems originating from under-sampling of rare interactions (i.e. missing links). They can easily accommodate qualitative or quantitative data, and can incorporate trait variation within species, such as values that vary along developmental stages or environmental gradients. We use three case studies to show that they can detect strong trait matching (e.g. predator-prey system), relaxed trait matching (e.g. herbivore-plant system) and barrier trait matching (e.g. plant-pollinator systems). Only by elucidating which species traits are important in each process, i.e. in determining interaction establishment, frequency, and efficiency, can we advance in explaining how species interact and the consequences for ecosystem functioning.

## Introduction

Species interactions form the architecture of biodiversity (Bascompte & Jordano 2007). There is growing recognition that community structure, stability and functioning depend not only on which species are present in a community, but also on how they interact (Tylianakis *et al.* 2008). Complex networks of biotic interactions such as predation, parasitism and mutualism provide essential information for conservation (Carvalheiro, Barbosa & Memmott 2008; Tylianakis *et al.* 2010), community stability and ecosystem functioning (Thompson *et al.* 2012; Peralta *et al.* 2014) and evolutionary processes (Jacquemyn *et al.* 2011; Fenster *et al.* 2015) that would be not possible from simple species occurrence data or analysis of pairwise interactions. Despite the growing literature describing species interaction networks, we still have a poor understanding on how this network structure comes to exist.

Disentangling what determines the occurrence of pairwise interactions, and at a higher level the structure of complex networks, is a key challenge for ecologists. Overcoming this challenge requires the identification of rules responsible for who interacts with whom. There is a great expectation that incorporating a trait-based approach can help us explain interaction occurrence among species. We refer here to traits in a broad sense, comprising characteristics that define organisms in terms of their ecological role, how they interact with the environment and with other species (Díaz & Cabido 2001). Recent studies indeed suggest that ecological interactions of all sorts could be described from the traits of the interacting species (Eklöf *et al.* 2013). The ability of these methods to predict the novel interactions following species invasions or following range shifts is, however, limited.

Traits are implicated in ecological dynamics at several concatenated levels of community organization (Fig. 1) and therefore could influence the occurrence of interactions in multiple ways. Some traits determine species distribution in a multi-dimensional environmental space and thus impact co-occurrence in space and time. Since the occurrence of an interaction requires the presence of the two species, traits involved in phenological matching or habitat filtering could constrain interactions. Life-history traits impact demography, abundance and biomass, thereby affecting the probability of encounter, a sine-qua-non prerequisite for two species to interact. Then, provided they encounter each other in space and time, the compatibility between traits of the two species (i.e. trait-matching constraints) will also determine whether or not they interact. Finally, the intensity and the impact of an interaction will determine the functioning of the network. How efficient a species is on a per capita basis is also likely to be mediated by its behavioural or physiological traits and how these match with those of the other species. Most work to date has focused on morphological trait matching and few, if any, has tackled several aspects at a time (see the review in Morales-Castilla et al. 2015). Our first objective here is to review what we know about each of these processes and assess their success and limitations to predict interactions. Our second objective is to propose a way forward to evaluate species trait matching, and how this can be integrated in the bigger picture, from species occurrences to ecosystem functioning.

**Figure 1:**
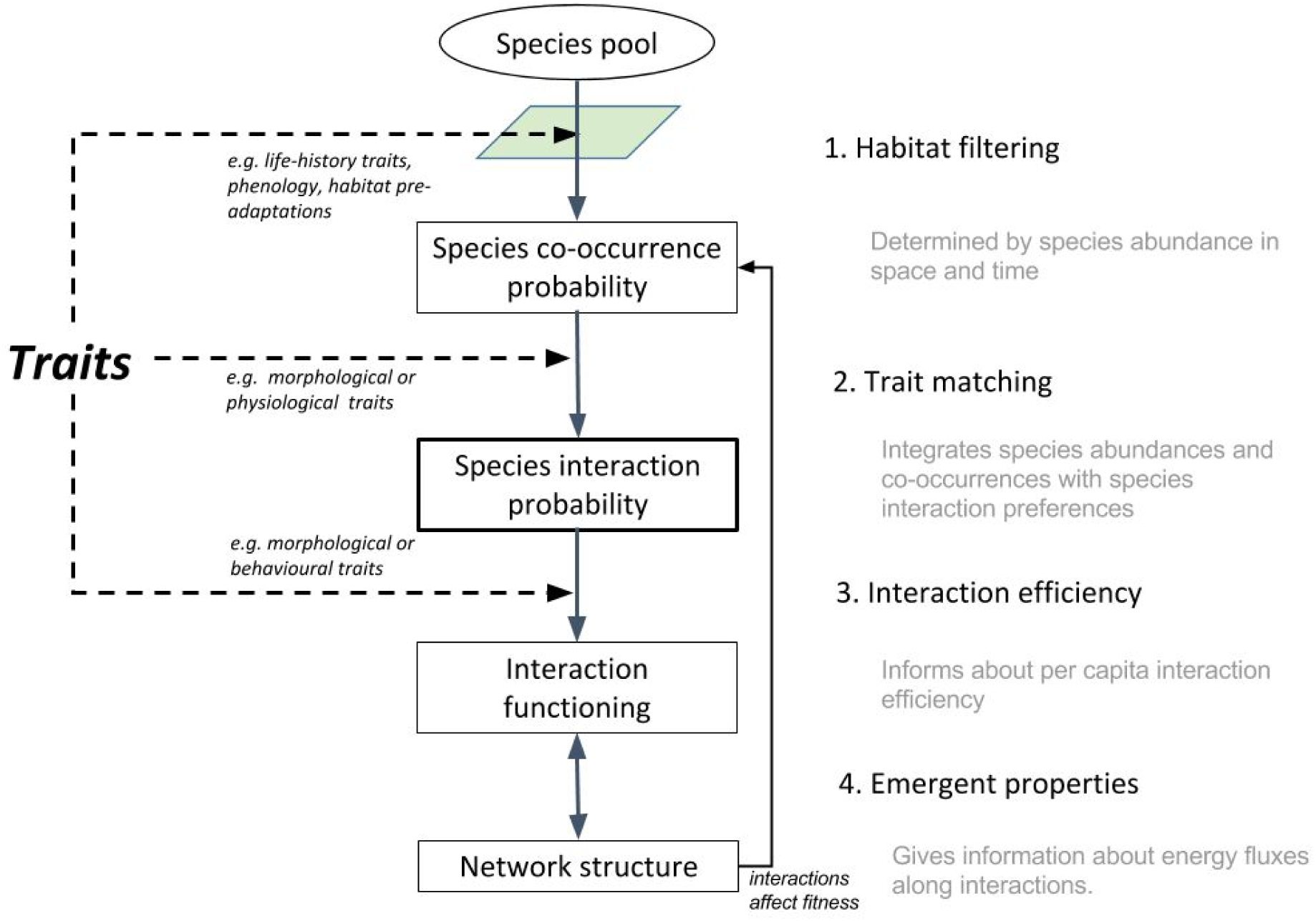
Scheme of how species traits can shape the interaction network at different stages.

## Traits governing species distribution and abundance in space and time

For two species to interact directly, the first requisite is that they co-occur in space. Given the heterogeneous distribution of most species, habitat filtering will constrain the pool of species co-occurring in a given region or microhabitat. Sharing ecological traits, like adaptations to particular environmental conditions, may hence be a prerequisite for two species to interact. Even in large, diffuse networks such as the global planktonic interactome, 18% of average community structure can be explained by environment alone, and these co-occurrences can be used successfully to predict interactions between taxa (Lima-Mendez *et al.* 2015). Microhabitat can have a strong influence for sessile organisms even within close proximity, as shown by interactions among mycorrhizas and plants, where rooting depth could preclude interactions between shallow rooted plants and fungi restricted to lower soil horizons. In fact, the concept of “habitat associations” as a driver of interactions has been pointed as the sole explanation for this interactions (Zobel & Öpik 2014), suggesting that both partners interact simply because they respond independently, but in the same way, to their environment.

Similar to species distribution in space, species co-occurrence will be determined by the synchrony of their activity periods at different temporal scales (i.e. daily, seasonal, interannual). In a network context, phenology has been widely used to explain forbidden links (Olesen *et al.* 2011; Encinas-Viso, Revilla & Etienne 2012; Olito & Fox 2015), that is, species present in the same communities that do not interact because they do not overlap in their seasonal activity periods. While phenology is usually studied as the timing when species are active during a season (e.g. plant flowering period), daily fluctuations of activity can also be important for defining when interactions among partners can occur. A clear example is the distinction between crepuscular vs. diurnal species (Herrera 2000), but more subtle activity fluctuations depending on daily temperature may be also relevant (Rader *et al.* 2013). In addition, some species may interact only with partners in a given life-history stage, for example, ectomycorrhizal fungi require hosts to be at least several years old and they do not interact with seedlings. This highlights the importance and complexity of the temporal aspects of interaction co-occurrence.

Co-occurrence is now explored as a mechanism driving interactions. Species turnover along ecological gradients is shown to be responsible for a large fraction of network variation in space (Poisot *et al.* 2012). This can be used to better understand the consequences of range shifts on the local food-web structure (Albouy *et al.* 2014). Alternatively, co-occurrence matrices could be also described with network metrics, provided that co-occurrence is constrained by interactions (Araújo *et al.* 2011). At smaller scales, phenological overlap during the season has also been used as a proxy for interaction probability (Bartomeus *et al.* 2013). More generally, we envision that species distribution models in combination with ecological and life-history traits (D’Amen *et al.* 2015) holds a promise to predict co-occurrence and potential interactions at multiple spatial scales and in response to global changes (Morales-Castilla *et al.* 2015).

Given that species co-occur in space and time, their abundance also determines the probability that two species interact (Canard *et al.* 2014). Abundant species are simply more likely to encounter each other than are rare ones. This mechanism has been called neutral because it does not rely on any niche differentiation. Thus, models that use species abundances to predict encounter probabilities have found that abundance alone can explain considerable variance in key aspects of network structure (Vázquez *et al.* 2007; Krishna *et al.* 2008; Olito & Fox 2015). Abundances are commonly used to develop a null model to reveal the added effect of trait-matching. However, life history traits, such as fecundity or longevity, will also constrain abundance. For plant communities, there is some consensus over which traits relate to abundance or dominance in the community, such as maximal height and position along the slow-fast continuum (e.g. leaf economic spectrum; Wright *et al.* 2004). Trait distributions over environmental gradients have been used to predict plant abundance and community structure (Shipley, Vile & Garnier 2006; Laughlin *et al.* 2012). Similarly, it is possible to relate life-history traits to animal abundances. For instance, species with fast life cycles (usually small, with high reproduction rates and short longevity) tend to be more abundant than large species with slow life histories (White *et al.* 2007). As a result, abundance is largely related to body size and position in the interaction network (Woodward *et al.* 2005).

## Trait matching

Trait matching between interacting partners has been identified for a variety of organisms. Plant corolla length and pollinator proboscis length is a classic example (Kritsky 1991), despite most pollination interactions now being considered to be quite generalized and hence little constrained by trait matching (Waser *et al.* 1996). Bird beak size and fruit size has also been shown to be tightly related to dispersal success (Galetti *et al.* 2013). In fishes, predator mouth gap and prey size are also strong determinants of predatory interactions (Cunha & Planas 1999). More complex relationships have been found for plants too, with specific leaf area changing plant-plant interactions from facilitation to competition, depending on resource availability (Gross *et al.* 2009). In general, more or less constrained matching mechanisms has been proposed for most interacting species ranging from arbuscular mycorrhizas and plants (Chagnon *et al.* 2013) to plants and herbivores (Deraison *et al.* 2015).

However, trait matching between individuals operate in addition to neutral processes to impact pairwise interactions. Despite advances in these respective fields (e.g. null model analysis: Vázquez, Chacoff & Cagnolo 2009; trait matching analysis: Dehling *et* al. 2014; Spitz, Ridoux & Brind’Amour 2014), we still lack a common analytical framework with which to evaluate the contribution of species traits to pairwise interactions, and at the higher level to the structure of interaction networks.

Isolating neutral effects from trait matching effects on network structure remains a challenge. While it has been shown that both neutral and trait-based null models can predict the general structure of interaction networks, such models are poor at predicting individual interactions (Vázquez *et al.* 2009; Olito & Fox 2015). This suggests that neutral and trait-based null models may predict the right structure for the wrong reason. One major problem that may preclude disentangling these processes is that traits could influence interactions directly via trait matching, or indirectly via environmental matching and co-occurrence. Hence, even if we succeed to partition the variance between neutral and trait matching components, this would ignore the fact that some of the ‘neutral’ variance was generated by species traits (as we outlined in the previous section). Thus, the influence of abundance versus traits can be seen as a path diagram where traits directly affect interactions and also affect abundances, which affect interactions (Fig. 1). We propose a framework that aims to integrate, rather than separate both processes.

A significant challenge before such an analysis can be achieved is to access completely sampled networks with which to validate models (Bartomeus 2013). While model evaluation requires a null prediction against which to compare empirical data, empirical network data have inherent uncertainties associated with the way in which they are sampled. Specifically, sampling completeness is rarely achieved when collecting interaction networks (Chacoff *et al.* 2012), and hence, some unobserved interactions may indeed occur (i.e. false absence of interactions). This would be less of a problem if the proportion of interactions that are sampled were constant, but this sampling efficiency can vary with local environmental conditions (Laliberté & Tylianakis 2012), species abundance and frequency, and of course, sampling effort. Thus, to truly understand the importance of trait matching for determining species interactions, the absence of an interaction in an empirical dataset cannot be used to infer true absence of that interaction in nature. The nature of the data therefore impedes the direct evaluation of probabilistic models (e.g. Rohr *et al.* 2010) and requires the development of methods based on observed interactions only.

Another challenge is that null models based on *a priori* rules for interactions have to be constructed using assumptions of which traits are critical for interaction establishment, and which trait values of interacting species constitute matching. Constructing and interpreting biologically meaningful null models that can tease apart or isolate the targeted process to be studied is not an easy task (Vázquez & Aizen 2003). As an alternative, recent attempts to understand trait matching directly from empirical data (Bartomeus 2013; Dehling *et al.* 2014; González-Castro *et al.* 2015) are promising, but they are still unable to integrate the relative contribution of neutral vs. trait-based process.

A final caveat to existing approaches is that most null models are constrained to use mean trait values at the species level, neglecting the variability among individuals of the same species. However, intraspecific trait variation, which can result from life-history stage, sexual dimorphism, or stochastic, environmental, genetic or epigenetic forces (Bolnick *et al.* 2011), has been shown to affect specific interactions such as competition, as well as overall ecological dynamics (González-Suárez & Revilla 2013).

To overcome these limitations, we model the probability of interaction among pairs of individuals given their traits, based on a framework developed by Gravel and colleagues (Gravel *et al.* 2013). The framework evaluates trait matching relationships while taking into account neutral constrains (Box 1), and using only information about observed interactions, thereby overcoming problems caused by under-sampling of rare interactions leading to false absences of interactions. The approach has been shown to be robust to incomplete network sampling. Several models, corresponding to different hypotheses, can be fit directly to raw data and accommodate complex trait matching response functions to both qualitative or quantitative interaction data. Finally, they can incorporate intraspecific trait variation, avoiding the loss of realism in species with trait values that vary along developmental stages or environmental gradients. In that way we provide a common toolbox with which to understand linkage rules across a variety of interaction types.

## Linking network structure to ecosystem functioning

The functioning of an ecosystem is driven not only by biodiversity (Cardinale *et al.* 2012), but also by the number, distribution, and efficiency of interactions (Duffy *et al.* 2009; Thompson *et al.* 2012). In fact, species interactions encapsulate most ecosystem process and functions (e.g. animal pollination, fruit dispersion) as well as energy fluxes (e.g. predation, parasitism), but network structure is still little explored for understanding ecosystem functioning (Thompson *et al.* 2012). A first step is to recognize that ecosystem functioning depends on how efficient interactions are among partners.

Interaction efficiency is the per capita strength of a single interaction link (Vázquez *et al.* 2015). Interaction efficiency is usually asymmetric among both partners, and can be positive for both partners (e.g., mutualism) or positive for one partner and negative for the other (e.g., parasitism, predation). Interaction efficiency can be driven by behavioural or morphological traits (e.g. large pollinators deposit more pollen, (Hoehn *et* al. 2008)) or by the extent of trait matching (e.g. pollinators with short tongues may be able to visit, but inefficiently pollinate long flower corollas). But empirical evidence measuring interaction efficiency is still scarce. It is interesting to note that interaction frequency will be strong predictor of ecosystem functioning only where interaction efficiency is relatively invariant in comparison to interaction frequency, as in pollination systems (Vázquez, Morris & Jordano 2005)

A second factor governing ecosystem function is overall network architecture. The field of biodiversity and ecosystem functioning has been largely developed for competitive interactions (Loreau 1998, 2010) but now theory (Thebault & Loreau 2003; Duffy *et al.* 2007) and empirical (Winfree *et al.* 2015) evidence shows it also expands to more complex communities. The concept of trophic complementarity for instance establishes that the distribution of interactions with resources and predators determines the relative importance of exploitative and apparent competition on ecosystem functioning (Poisot, Mouquet & Gravel 2013). Ecosystem functioning will be promoted by low overlap among species of a given level in both their interactions with the resources and the predators (Peralta *et al.* 2014). Given the role of traits in determining these interactions, functional diversity and identity will contribute positively to ecosystem functioning (Cadotte, Carscadden & Mirotchnick 2011; Gagic *et al.* 2015). These emergent properties of the network could be easily incorporated into our framework, via a feedback from network structure to the function determined by individual interactions in Fig. 1.

## Case studies

We re-analyzed three datasets on different systems ranging from antagonistic to mutualistic interactions to illustrate the utility of our framework (which is presented in Box 1). We first describe in Box 1 a situation where trait matching is a strong driver of interaction composition (because of a strong predator-prey body size relationship for marine fishes). The parameters of the fitted model can subsequently be used for predicting interactions among species that currently do not co-occur but may do so in the future, for example as a consequence of range shifts under climate change (Albouy *et al.* 2014) or species invasions. The experimental data (Deraison *et al.* 2015) on the relationship between grasshoppers incisive strength and leaf dry matter content alternatively shows weak trait-matching when using binary data (Fig 2A). However, weighting interactions by their consumption rate frequency removes bias in parameter estimates, and the model consequently exhibits clearly that strong-mandibled grasshoppers prefer plants with higher content of dry matter, as reported in the original paper (Fig 2B). Interestingly, in this example, the trait matching function directly maps on the per capita efficiency (i.e. consumption rate).

**Figure 2:**
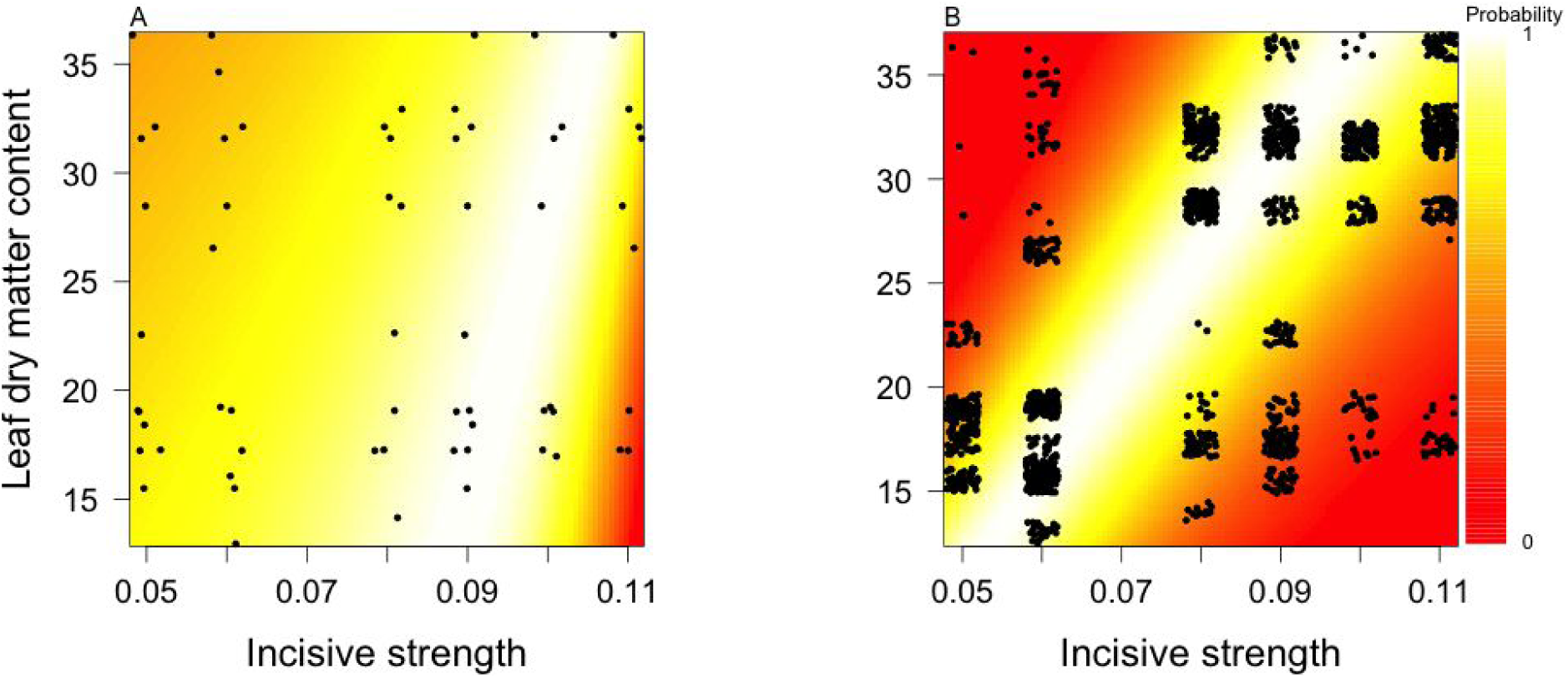
Model for grasshopper and plant interactions unweighted (A) and weighted (B) by frequency of interaction. The probability of interaction between a grasshopper and a plant follows a positive relationship between incisive strength and plant leaf dry matter content. Note that the overlapping data in B have been jittered to appreciate the different frequencies of particular interactions. The likelihood for (A) is similar to the neutral model, while much better in (B), indicating that the frequency of interactions must be taken into account to better reveal the trait-matching constraint.

The previous example used trait mean values at the species level, treating all individuals of the same species as identical, and ignores individual variability. However, within-species variability has been shown to be very relevant for ecological dynamics. Our framework also allows the capture of inter-individual differences when evaluating parameters of trait-matching functions. Both the predator-prey model shown in Box 1 and the example of Fig. 3 use individual specimen measurements of traits. For the latter, we measured each pollinator individual in (Bartomeus, Vilà & Santamaría 2008) and related pollinator tongue length (measured using the allometric relationship between tongue length and body inter-tegular span within each bee family; Cariveau *et al.* 2015) with plant nectar holder depth. Average measures were used for plants. This example illustrates how individual variability can be accommodated by our framework. In addition, this model uses independent information to describe the trait abundance distribution of plant species. In the past examples, abundance was inferred from the network of interactions, but in this case, independent transect measures of percent plant cover in the site are available (Bartomeus *et al.* 2008), and were used to weight the trait distribution of plant traits that the pollinators are selecting. We show that an interesting trait-barrier model emerges, where small-tongued individuals cannot access deep flowers, but long-tongued species can access both deep and shallow flowers (Fig. 3). However, in this case, the neutral model provides a very similar likelihood to the integrated model, indicating that under such weak constraints (most pollinators can access most plants), abundance is the main determinant of interaction probability.

**Figure 3:**
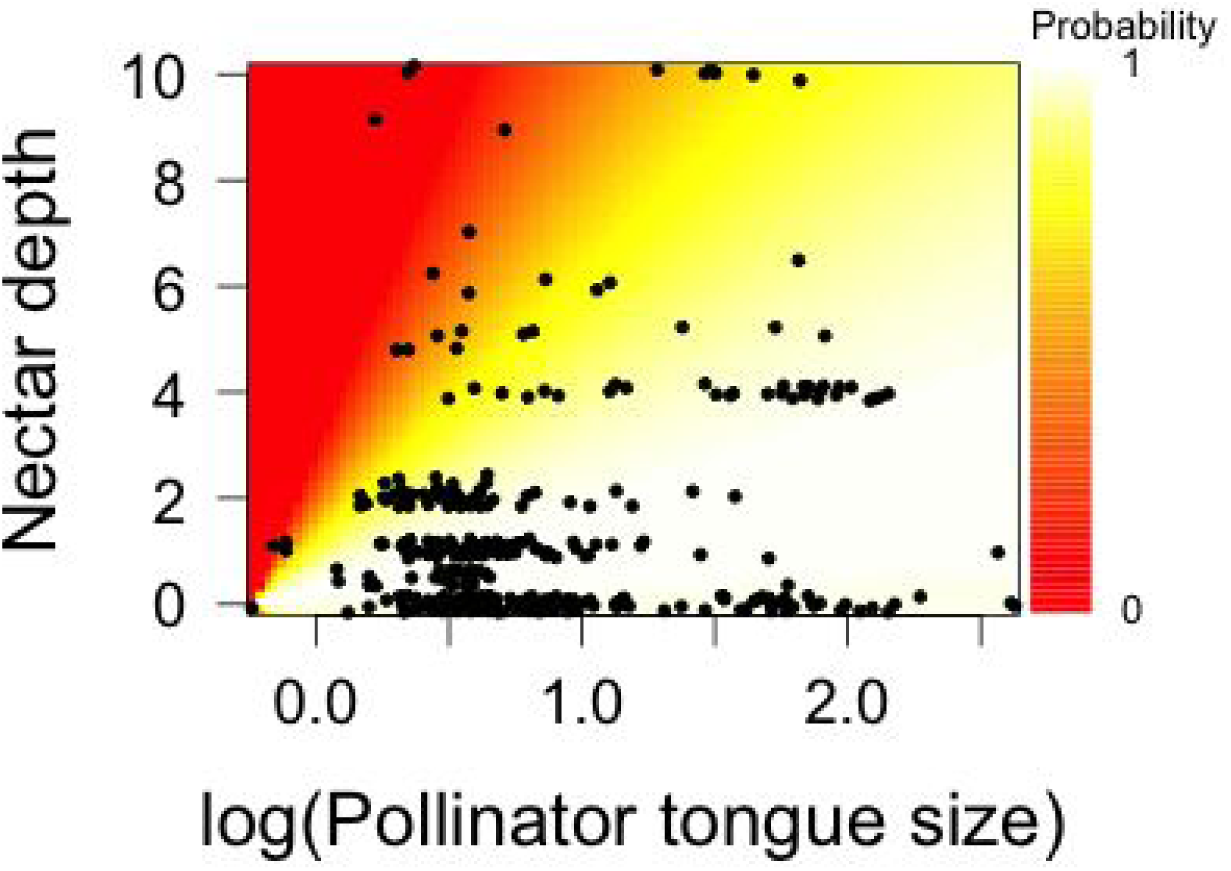
Model for plant-pollinator interactions weighted by the frequency of interactions. We show that only a few interactions among small tongue sized bees and long corolla depth flowers are not realized (red area), while the rest of interactions are explained mainly by neutral process.

A detailed guide to running all these models and interpreting the output can be found here (https://github.com/ibartomeus/trait_match).

## Discussion and conclusions

Quantifying the trait matching relationships across species may help us to understand how networks are structured. For example, the nested structure of plant-pollinator networks may be driven from species abundance (Vázquez *et al.* 2009) or from barriers to certain interactions (Stang, Klinkhamer & van der Meijden 2006). In contrast, the strong trait matching observed in plant-herbivore interactions (e.g. plant defenses limiting herbivory for all but a few tolerant species) can produce more modular networks where interactions depart more from the null expectation based solely on abundance (Thébault & Fontaine 2010). Even within plant-pollinator interactions, bird-plant networks are more specialized than insect-plant networks, which is also reflected in their degree of trait matching (Maglianesi, Böhning-Gaese & Schleuning 2015). Our framework is however limited to pairwise interactions and future work will have to investigate how the trait distributions in a community constrain the emergent network properties. Moreover, trait-matching constraints describe potential interactions, but may not always reflect realized interactions. (Poisot, Stouffer & Gravel 2015) nicely develop this concept by adding to the neutral and trait matching components an error term that represents the emerging properties of the network. Our models also have an associated error that allows explicit recognition of this uncertainty regarding the outcome of predicted interactions.

Parameterized trait matching functions not only provide a better understanding of the drivers of interactions, but they also allow prediction of novel interactions following deliberate introductions (e.g. of crop species or biological control agents) or unintentional invasions and range shifts (Morales-Castilla *et al.* 2015). Proxies of trait similarity, like phylogenetic distance, have already been successfully used to predict interactions of exotic species (Pearse & Altermatt 2013) and adding traits has the potential to enhance this approach. Species losses and gains following global changes are threatening most ecosystems, and it is simply impossible to measure all potential interactions in the field. Tools are consequently required to assess how the interaction network will rewire. We know that exotic species invading a community get easily integrated into the recipient network of interactions (Albrecht *et al.* 2014), and that after species turnover in a community, the remaining species reshuffle their interactions to adjust to the new composition (Kaiser-Bunbury *et al.* 2010). However, our predictive power in these situations is still limited.

Careful selection of the right set of traits is, however, a critical step. We have seen that traits constraining interactions could potentially comprise all morphological and physiological species characteristics, and hence, are quite specific for each interaction type. A good *a priori* knowledge on the biology of the species and type of interaction involved is needed to select the right trait combinations. For example, we also explored whether body-size drives host-parasite relationships using the (Tylianakis, Tscharntke & Lewis 2007) dataset, but in this case all models performed poorly because body size appears less important for determining each of these interactions. In this case, body size does not constrain host and parasite interactions because the largest parasitoid is smaller than the smallest host, which allows all types of body size combinations.

Alternatively, spurious trait matches could be found when some traits are correlated. For instance, traits like body size correlate allometrically with several other morphological traits (Woodward *et al.* 2005) and might therefore provide a wrong causal explanation of the interactions. One strong limitation for some interactions, such as fungi and plants, is that the traits governing interactions remain somewhat unclear (Tedersoo *et al.* 2008; Martínez-García *et al.* 2015). The challenge for the future will be to determine and quantify the actual traits governing these interactions, including their variability among individuals or genets.

In conclusion, different traits can inform us about how species form networks of interactions. For some interaction types, like mycorrhizal fungal interactions, traits affecting species co-occurrence can be the most relevant for understanding the occurrence of interactions. Conversely, for other interaction types, like those between predators and prey, morphological and physiological traits may be the main determinants of who interacts with whom. Understanding which mechanisms are driving pairwise interactions is key to predicting how communities will respond to global change. Interactions regulated by co-occurrence will be more likely to be affected by climate change (e.g. changing phenologies and distributions), while changes in dominance following disturbance may redistribute the interactions in neutral-driven networks. Non-random species extinctions are also expected to affect more drastically interactions regulated by strong trait matching (Larsen, Williams & Kremen 2005). Even though the unknowns are still too great to draw general conclusions about how communities are structured and what implications this has for ecosystem functioning, we are now armed with an increasing number of empirical examples and the right analytical tools to move beyond descriptive patterns and into predictive analysis that allows us to tackle important questions regarding community assembly in the face of global change.

## Box 1.1 A bayesian method for evaluating trait matching as a determinant of network structure

We are interested in evaluating from empirical data a function describing the probability of an interaction between species *i* and *j* based on their respective sets of traits *T*_*i*_ and *T _j_*. Building upon the example developed by (Gravel *et al.* 2013) *we aim to fit a statistical model that will relate the probability with which an interaction occurs to the set of traits of the two species:*

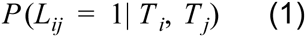

Which reads as the probability of observing an interaction *L* between species *i* and *j* given the traits *T _i_* and *T _j_*. The function describing this probability could take any form. For the sake of the example here, we will consider a gaussian function (i.e. a function that assumes a linear relationship between *T _i_* and *T _j_*) to represent the interaction niche ((Williams, Anandanadesan & Purves 2010), see below) but other functions, such as a high order polynomial or even regression trees, could be considered as well. The gaussian function is however convenient because it is easy to integrate.

Equation 1 could be fit directly to empirical data. To do so, the required data should contain information of presence and absence of interactions (e.g. (Rohr *et al.* 2010). The problem we are facing, however, is that records of the true absence of interactions are often not available in most datasets of ecological interactions, and when available, there might be considerable uncertainty in these absences (i.e. false negatives due to insufficient sampling). We therefore present a bayesian methodology to fit Eq. 1 indirectly, using only information about the observed interactions. Presence only data have information about the traits of species *i*, of species *j* and the interaction *L*_*ij*_ = 1. We consequently revise the problem and model the probability of sampling trait *T _i_*:

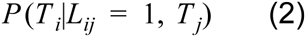

Which could be interpreted as the probability that we pick trait *T _i_* from the trait distribution we model, given we know there is an interaction between species *i* and *j* and the trait *T _j_*. This equation provides the likelihood for any observation of an interaction based on traits of the two species. We now use Bayes’ theorem *p*(*A*|*B*)*p*(*B*) = *p*(*B*|*A*)*p*(*A*) to decompose Eq. 2, yielding the following posterior distribution of prey size, given the predator size:

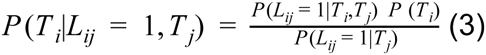

*P* (*T*_*i*_) is the probability density function for the trait *T _i_*. It corresponds to the probability of this trait in the regional pool. It could thus be weighted by abundance because the most abundant species are more likely to be sampled. The denominator is the marginal distribution of the interaction probability, computed as the integral of the numerator over the whole distribution of the trait *T _i_*:

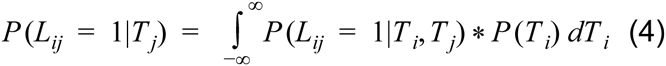

The overall principle of the method is best interpreted in light of the below figure. Take for instance the case of a predatory fish species of size *M*_*pred*_ selecting prey from the entire distribution of body mass of a set of prey, *P* (*Mprey*). We know that larger fish typically feed on smaller ones because they must catch and handle the prey with their mouth. The frequency distribution of prey size will indeed influence the distribution of the body mass in the diet of the predator. A predator will tend to feed most often on the most abundant prey, which is a neutral component to the interaction probability. The predator does not select from that distribution randomly, however, but rather it targets only a specific range (given by Eq.1 the niche component). Both the available prey distribution, *P* (*Mprey*), and the posterior prey distribution, *P* (*Mprey*|*L*, *M*_*pred*_), are illustrated in panel B. The posterior prey distribution will be somewhere between the regional prey distribution and its niche. The model therefore integrates both neutral and trait-matching constraints.

As a side product, the denominator informs us of the generality of the consumer. This integral might be tricky to compute analytically, depending on the form of Eq. 1 and the type of distribution, but most software nowadays offer easy ways to compute it numerically. Here the usage of the linear function simplifies computations. We provide an example in panel C for trophic interactions in marine systems. We use data from (Barnes *et al.* 2008), using log body size for the predator (*M*_*pred*_) and prey (*Mprey*). The trait-matching function is based on the niche model (Williams & Martinez 2000), where the main niche axis is the log of body size. In this situation, both *T*_1_ and *T* _2_ are the same trait. The log body size of the predator determines its optimum and the range of its niche, while the log prey size its niche position (Gravel *et al.* 2013). We consider a gaussian function to represent the probability of an interaction given the size of the predator and the prey:

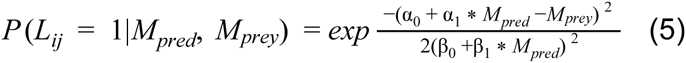

Where α_0_, α_1_, β_0_ and β_1_ are fitted parameters describing the linear relationship between the predator size, its optimum (α_0_, α_1_: intercept and slope) and the range (β_0_, β_1_) of its niche (other shapes could be used as well). One tricky issue might be to gather information about the trait distribution. We assume here that the distribution of the data provides an adequate representation of the distribution of potential prey sizes. However, one could in principle take the average and the standard deviation of the trait distribution at the species level or at the individual level. We consider a normal distribution for the log of prey body size.

### Pure neutral interactions

The above model integrates both trait-matching and abundance constraints. The model could be simplified to account only for the effect of abundance (trait distributions) to reveal the importance of the trait-matching constraint. A neutral model in this framework is found when an interaction is equally probable, irrespective of the traits of the two species involved in the interaction. In other words, *P* (*L*_ij_ = 1|*T*_*i*_, *T*_*j*_) = *k*. In this situation, the probability of sampling trait *T*_*i*_ is given by:

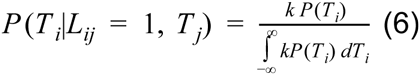

Which reduces to the distribution *P* (*T*_*i*_) because the integral of the posterior distribution function sums to 1 by definition, and the constant k can be simplified.

### Pure niche interactions

Alternatively, one could want to compare to the situation where interactions are purely determined by trait-matching constraints. In this situation, we consider the distribution of the trait *T _i_* to be uniform within the range of the observed traits. The probability of sampling trait *T*_*i*_ in this situation is by definition 1/(*max*(*T*_*i*_) − *min*(*T*_*i*_)) for all trait values. The denominator simplifies to the integral of the gaussian function (Eq. 5), 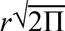, and thus the posterior distribution of the pure niche model takes the form:

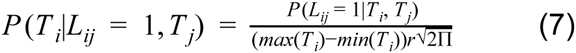

### Multi-trait expansion

The extension to multiple trait matching constraints is straightforward to perform. Each species has multiple traits, denoted *T*_*i*1_, *T*_*i*_2… This extension requires an assumption that the different trait matching functions are independent (cases of non-independence are possible but beyond the scope of the current paper). Because of this assumption, the joint probability is easily expanded using the relationship *P* (*x*, *y*) = *P* (*x*)*P* (*y*):

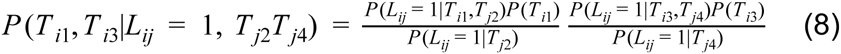

All of the R code necessary to perform this analysis is provided here: https://github.com/ibartomeus/trait_match.

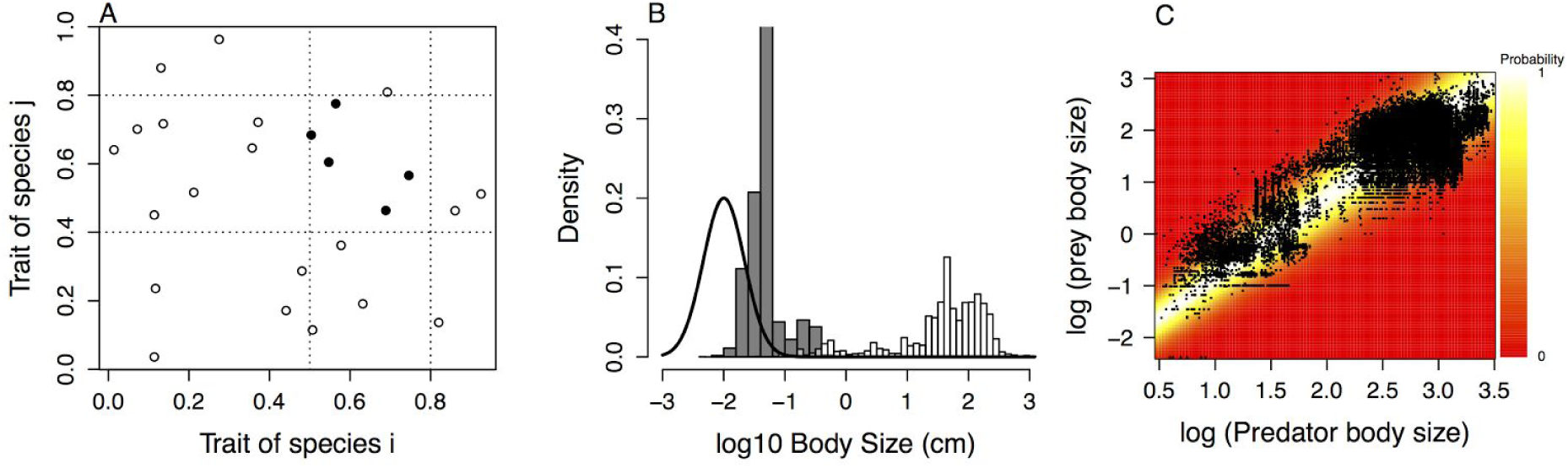

Illustration of the quantitative framework to evaluate a trait-matching function. A) Conceptual representation of a trait-matching constraint. Interactions (in black) are observed only when both species have traits that are compatible. However, we often do not have information on the white dots (species that are present but do not interact). B) Representation of the density function for available body size in the (Barnes *et al.* 2008) dataset, the trait-matching function (black line) and the observed distribution of prey size for a given predator (black bars). C) Representation of the observed interactions (black dots) and the maximum likelihood estimate of the trait-matching function (from low probability in red to high probability in yellow).

